# Pax6 homologs are required for patterning both visual systems of the daddy-longlegs *Phalangium opilio*

**DOI:** 10.64898/2026.04.03.716372

**Authors:** Ethan M. Laumer, Sophie M. Neu, Benjamin C. Klementz, Pranil Panda, Emily V.W. Setton, Prashant P. Sharma

**Affiliations:** University of Wisconsin-Madison, Department of Integrative Biology, Madison 53706, Wisconsin, United States; The Whitney Laboratory for Marine Bioscience, Department of Biology, University of Florida, St. Augustine, FL, 32080, USA

**Keywords:** morphogenesis, eye development, paralogy, vestigial organs, retinal determination network

## Abstract

The evolution of visual systems has compelled numerous investigations of developmental processes underlying eye patterning across Bilateria. It is well-established that homologs of the transcription factor Pax6 play a highly conserved role in eye fate specification and are at the top of the retinal determination gene network (RDGN) hierarchy. In insects, the two Pax6 homologs *eyeless* (*ey*) and *twin of eyeless* (*toy*) are required for the development of the two visual systems broadly found within the phylum (i.e., median and lateral eyes). Curiously, Pax6 homologs do not appear to maintain this function in well-studied chelicerate models, with emphasis on spiders, a lineage of arachnids with great diversity of eye form and acuity. It was recently proposed that the gene Pax2 (*shaven*; *sv*) may have subsumed the role of eye fate specification in chelicerates, a hypothesis predicated upon the observation that one of two spider Pax2 copies is strongly expressed in the developing lateral eyes during embryogenesis. However, no functional data are available for any Pax homologs across Chelicerata. We examined the incidence of Pax family genes across Chelicerata, as well as interrogated the expression and function of Pax2 and Pax6 homologs in the daddy-longlegs *Phalangium opilio*, an arachnid recently discovered to bear a highly plesiomorphic arrangement of visual systems. Here, we show that *ey* and *toy* are expressed early in the developing head lobes of *P. opilio*, whereas *sv* is not expressed until well after stages when downstream RDGN members (*eyes absent* and *sine oculis*) are already activated. Gene silencing of *ey*, *toy*, and *sv* individually had no discernible effect on eye development. By contrast, double knockdown of *ey* and *toy* resulted in an array of median eye defects, spanning loss of some cells of the eye to total loss of the median eyes. Gene expression assays also showed that depletion of the two Pax6 copies resulted in failure of the vestigial median and vestigial lateral eyes. These data are consistent with a conserved role for Pax6 homologs in patterning both visual systems and all three eye pairs in the daddy-longlegs. Our results comprise the first functional data for Pax6 genes in any chelicerate and suggest that heterochronic shifts in expression, rather than changes in function, underlie the atypical dynamics of Pax genes in derived arachnid groups such as spiders.

## Introduction

Eye fate specification is an intricate process that is essential for initiating the development of this complex organ. Across Bilateria, homologs of the transcription factor Pax6 play a central role in specifying eye fate, suggesting deep evolutionary conservation of the eye patterning program [1]. Broadly, the Pax family of transcription factors is characterized by the presence of a conserved 128-amino acid domain (PRD), which in turn is comprised of two helix-turn-helix (HTH) DNA-binding subdomains PAI and RED [2]. Abrogation of Pax6 function in mammalian models results in failure of eye formation and brain morphogenetic defects [3,4]. Similarly, loss-of-function experiments for arthropod homologs of Pax6 in insect models elicit defects of the two visual systems (ocelli or median eyes; and compound or lateral eyes), as well as head defects [2,5–8]. The functional significance of Pax6 sequence conservation is underpinned by seminal experiments showing that exogeneous Pax6 coding regions from a diversity of species can induce ectopic initiation of eye development in both the fruit fly *Drosophila melanogaster* and the clawed frog *Xenopus laevis* [9–11]. The accumulation of expression and functional data for Pax6 homologs in different metazoan clades has broadly supported their central role in eye morphogenesis [7,12–14], a function that may predate the origin of Bilateria [15]. Recent work in the fruit fly has also suggested that the partitioning of the two visual systems of arthropods is associated with differential expression levels of the two arthropod Pax6 homologs, *eyeless* (*ey*) and *twin-of-eyeless* (*toy*), with *ey* playing a more prominent role in the patterning of the compound (lateral) eyes and *toy* of the ocelli (median eyes) [16]. Both paralogs play synergistic roles in patterning the lateral eyes, whereas *toy* has a more prominent role in patterning the median eyes. Altering the relative expression of these two homologs can lead to a spectrum of eye defects, whereas altering the expression of downstream targets such as *eyes absent* (*eya*) can induce homeotic transformation of ocelli to compound eyes [16]. While functional datasets from other insect and crustacean models are not as experimentally advanced, these broadly support the conservation of the retinal determination gene network (RDGN) in *D. melanogaster*, with emphasis on the prominent role of Pax6 genes in eye morphogenesis [7,8,13].

An odd exception to this understanding occurs in Chelicerata, the sister group of the remaining arthropods. Like in insects, two visual systems (median and lateral) occur in chelicerates, though one or both are reduced or completely lost in certain chelicerate orders (e.g., Pycnogonida; Solifugae; Ricinulei; Palpigradi) [17]. The best studied visual systems in chelicerates are those of horseshoe crabs and spiders, which differ markedly from each other [18,19] (Fig 1). The most prominent eyes of horseshoe crabs are the lateral facetted eyes, which bear over 1000 ommatidia and are adapted for changes to vision between day and night [20,21]. Given the early appearance of facetted eyes in Cambrian arthropods, the structure and organization of the horseshoe crab facetted eye has featured prominently in reconstructing the evolution of arthropod vision [21,22]. Posterior to the lateral faceted eye is a second rudimentary lateral eye that is formed during early development in the trilobite larva. Additionally, a trio of eyes occurs in the dorsal midline, comprising a pair of median eyes (ocelli) and an endoparietal eye, formed by fusion of a larval median eye pair; the median eyes play important roles in detection of UV light and the establishment of lunar cycles. A pair of ventral eyes is located near the mouth, but their function is unknown [21]. By contrast, spiders bear a pair of median eyes and (typically) three pairs of disaggregated lateral eyes, though the spatial arrangement and number of eyes varies markedly between families [19,23]. The lateral eyes are distinguished from the median eyes by the presence of a reflective crystalline layer called the tapetum, as well as the arrangement of the rhabdoms with respect to the photoreceptor cell bodies (inverted in the lateral eyes with respect to the median eyes) [19]. In spiders with highly developed vision (e.g., Salticidae; Lycosidae), visual acuity is typically associated with the median eyes and the anterior pair of lateral eyes [24].

**Fig. 1.**
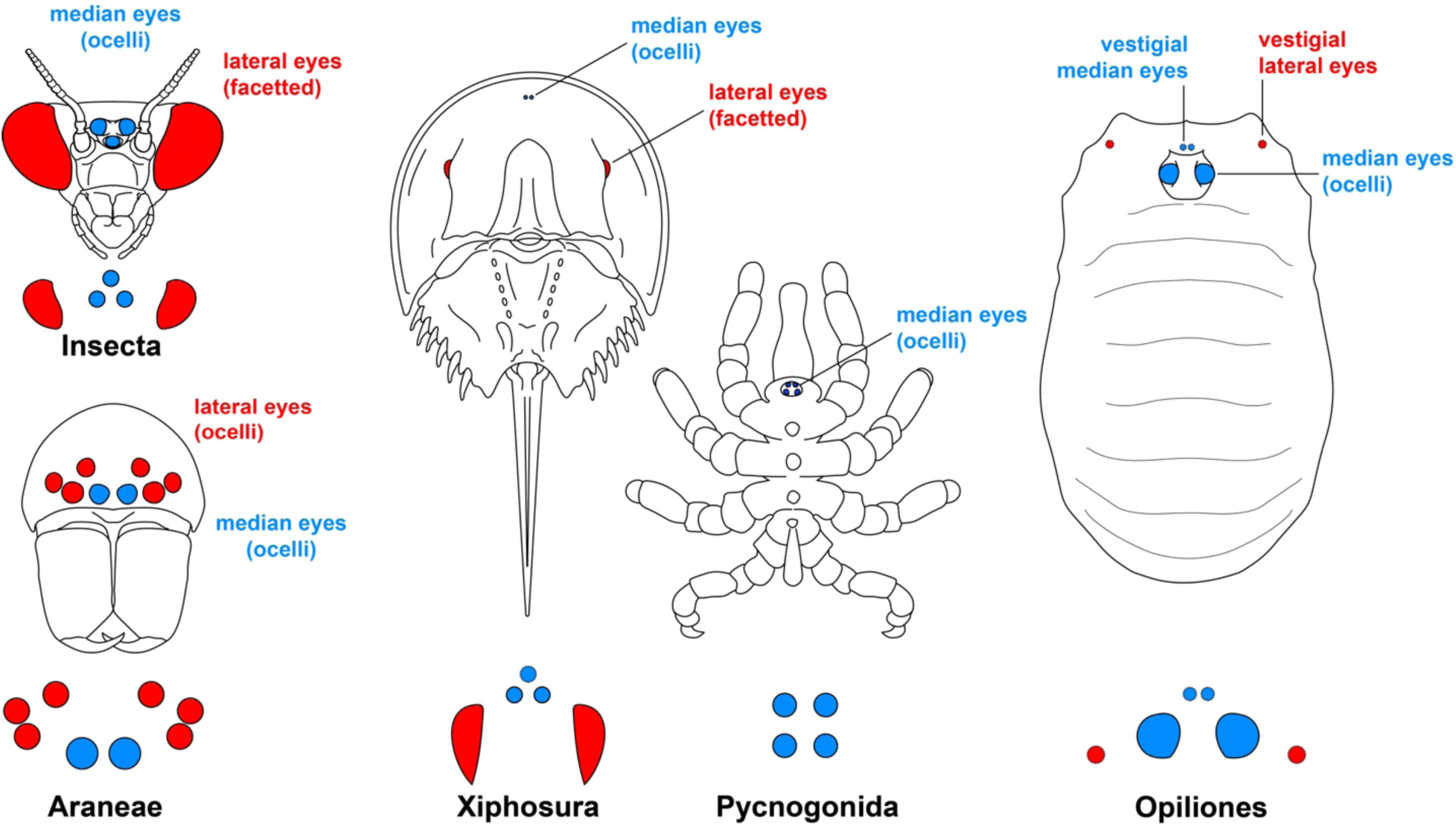
Organization of arthropod visual systems, with emphasis on Chelicerata. Red icons denote homologs of the lateral visual system, which includes the facetted eyes of insects and horseshoe crabs (Xiphosura), as well as the disaggregated ocelli of spider lateral eyes (Araneae). Blue icons denote homologs of the median visual system. Diagrams below line drawings represent simplified schematics of visual systems in each lineage. Note the incidence of two pairs of median eyes in sea spiders (Pycnogonida), as well as Opiliones (one pair vestigial). In Xiphosura, the third median eye shown in the schematic is the result of the fusion of paired larval median eyes. Additional eyes of Xiphosura are not shown for simplicity.

Despite these structural and functional differences, horseshoe crabs and spiders are united by the absence of Pax6 expression in the eye primordia during embryonic stages when eye fate specification is thought to occur. Embryonic expression data are limited for horseshoe crabs, but a previous survey of one of the five Pax6 homologs (the result of lineage-specific whole genome duplications) that occurs in *Limulus polyphemus* found no evidence that *Pax6* was expressed in the eye primordia [25]. Comparative data are more readily available for several spider species, including exemplars of different degrees of visual acuity. Initially, it was thought that the Pax6 homologs *ey* and *toy* were altogether absent from developing eye primordia, suggesting a Pax6-independent mechanism underlying eye fate specification across chelicerates [26,27]. However, subsequent explorations of earlier stages of development revealed that both copies were expressed in a broad domain demarcating the anterior head in the spider *Parasteatoda tepidariorum*, with subsequent diminution of transcription by the stages where the expression of downstream RDGN target genes *eyes absent* (*eya*) and *sine oculis* (*so*) is initiated [28]. These expression patterns were later corroborated in several other spider species [29]. It was therefore postulated that *ey* and/or *toy* may serve the function of establishing eye precursors indirectly and relatively early in development through the activation of *eya* and *so*, but are not required for the maintenance of these downstream targets [17,28]. While this explanation is plausible, no functional data exist supporting any role for Pax6 homologs in spider eye development, due to a combination of inefficiency of RNAi in this species [30–32] and the lack of reproducible gene editing tools in spiders [33,34]. In addition, it was recently shown that the same early expression domains of *ey* and *toy* occur in the anterior margin of the head in the blind oribatid mite *Archegozetes longisetosus*, suggesting that *ey* and *toy* plausibly retain a conserved role in head patterning as previously reported for insects [35].

An entirely separate hypothesis for the patterning of chelicerate eyes is the recent proposition that Pax2 (*shaven*; *sv*) may have taken on the function of eye fate specification in arachnids [36]. Two copies of *sv* (distinguished as *Pax2.1* and *Pax2.2*) occur in spider genomes, owing to an ancient genome duplication in the common ancestor of spider and scorpions (Arachnopulmonata) [37–39]. In embryos of two spider species, it was found that *Pax2.1* was expressed in the lateral eye primordia of two spider species, whereas *Pax2.2* was expressed elsewhere in the brain and in developing sense organs throughout the body [36]. Based upon these gene expression patterns, the authors proposed an intriguing model wherein Pax2 replaced at least the lateral eye fate specification function of Pax6 in spiders (*Pax2.1* is not associated with the median eyes), if not all chelicerates.

The inference that *Pax2.1* alone represents a key player in chelicerate lateral eye fate specification was later undermined by the observation that both *sv* ohnologs are expressed in both the median and the lateral eyes of the scorpion *Centruroides sculpturatus* [40]. Moreover, the single-copy homolog of *sv* in the daddy-longlegs *Phalangium opilio* exhibits an identical expression pattern to the two scorpion paralogs [40], suggesting that the restriction of spider *Pax2.1* to the lateral eyes represents a spider-specific phenomenon and cannot be generalized to all arachnids [41]. More generally, given that whole genome duplications are not characteristic of all chelicerates, the dynamics of duplicated genes in groups like spiders and horseshoe crabs may not be representative of the entire clade. Still, the role of *sv* (like Pax6 homologs) in arachnid eye development remains entirely untested because no functional data exist supporting either hypothesis.

An investigation of Pax gene function in chelicerate eye development would best be served by investigating a model with (1) both visual systems, (2) single-copy homologs of RDGN genes (due to the possibility of functional redundancy incurred by whole genome duplications), and (3) the availability of robust functional tools. The single best candidate for such an investigation is the daddy-longlegs *P. opilio* [42–44]. Though it was long thought that extant Opiliones only bear a single pair of eyes, we recently showed that three pairs of eyes develop in embryos of *P. opilio* (median eyes, vestigial median eyes, and vestigial lateral eyes), with two pairs of these persisting into adulthood (median and vestigial lateral pairs) [40]. Opiliones therefore exhibit a more plesiomorphic organization of visual systems than either spiders or scorpions, and more closely resemble the horseshoe crab and pancrustacean visual system arrangement.

Here, we investigated the function of *sv* and both Pax6 homologs (*ey* and *toy*) in developing embryos of *P. opilio* using RNA interference (RNAi). We show that Pax6 homologs are expressed in early stages, consistent with the hypothesized role in early eye fate specification, whereas *sv* is not expressed until after the initiation of *so* and *eya* expression. We also show that double knockdown of *ey* and *toy* results in defects of the eyes and loss of opsin expression in all three eye pairs, whereas *sv* RNAi has no effect on any eye type, paralleling experimental results in insect models. These first functional datapoints for Pax genes in Chelicerata suggest retention of a deeply conserved role for Pax6 genes in the specification of both arthropod median and lateral visual systems.

## Results

### *ey* and *toy* are retained across Arthropoda

A recent survey of Pax genes in insects reported some ambiguity in the presence of both *ey* and *toy* in chelicerates and myriapods, the two taxa subtending Pancrustacea in arthropod phylogeny [8]. The challenging orthology of chelicerate Pax6 genes is thought to result from the lack of two diagnostic sites in the Pax6 sequences of chelicerates (these two sites are understood to be specific to Pancrustacea), the paucity of information content in multiple sequence alignments restricted to the PRD domain, and limitations in taxon sampling [2]. We therefore performed a survey of the Pax family sampling 11 out of the 14 orders of Chelicerata. Broadly, the number of Pax homologs per species varied more in chelicerates than previously reported in insects, a result consistent with multiple waves of whole genome duplication in two separate branches of the chelicerate tree of life (Fig 2A). The largest complement of Pax genes was found in the horseshoe crab *Carcinoscorpius rotundicauda* (n = 33), whereas the smallest was found in the pseudoscorpion *Cordylochernes scorpioides* (n = 8).

**Fig. 2.**
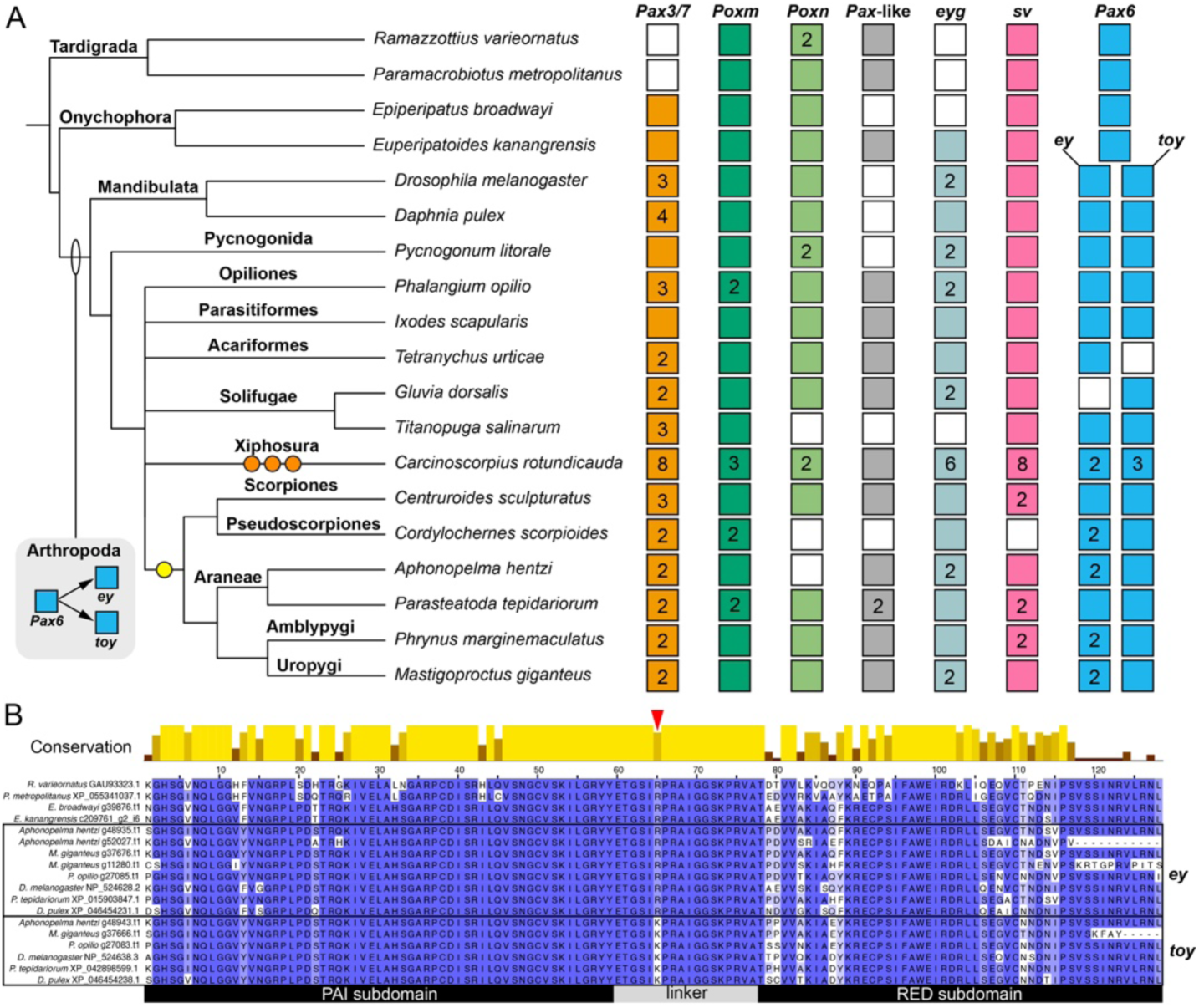
Pax gene complements across Chelicerata reveal lineage specific patterns of paralog retention in Arachnopulmonata. (**A**) Distribution of Pax homologs mapped onto a simplified phylogeny of panarthropods. Note the ancestral duplication of Pax6 in the common ancestor of Arthropoda. Incidence of Pax homologs are indicated by colored boxes (following Gutiérrez Ramos and Pick [8]); numbers inside boxes denote the number of gene copies. Yellow and orange circles represent whole genome duplication events. (**B**) Multiple sequence alignment of the PRD domain for selected exemplars of Panarthropoda, indicating diagnostic site for *ey* and *toy* in the linker region between the PAI and RED subdomains (red arrowhead). Design for panel (**B**) follows Friedrich [2].

In a phylogenetic analysis inferred from multiple sequence alignment of the PRD domain, two Pax6 homologs were recovered in all arthropods except *Tetranychus urticae* (Acariformes; *ey* only), *Gluvia dorsalis* (Solifugae; *toy* only), and *Phrynus marginemaculatus* (two copies of *toy* only). As previously noted, the incidence and expression of *ey* and *toy* have been described in the acariform mite *A. longisetosus*, suggesting that the absence of one copy in *T. urticae* may be a lineage-specific loss. Similarly, the lack of *ey* in *G. dorsalis* is likely attributable to the incompleteness of this genome; both *ey* and *toy* were found in the developmental transcriptome of a second Solifugae species (*T. salinarum*). By contrast to arthropods, only one Pax6 copy was found in both Onychophora and Tardigrada genomes, supporting the inference that *ey* and *toy* are the result of an arthropod-specific duplication [2].

As observed previously, the relationships between *ey* and *toy* are challenging to decipher using phylogenetic analysis of the PRD domain alignment alone, which did not result in mutually monophyletic clusters corresponding to *ey* and *toy*. To augment information content, we performed a separate analysis of only Pax6 homologs using full length sequences, which was able to resolve *ey* and *toy* as mutually monophyletic clusters (S1 and S2 Fig). Comparison of sequence data also corroborated the utility of a diagnostic site in the linker between the PAI and RED subdomains that reliably distinguishes *ey* from *toy* (Fig 2B).

The recent survey of Pax genes in insects also reported some ambiguity in the homology of Pax3/7 homologs (in *D. melanogaster*: *gooseberry*, *gooseberry-neuro*, and *paired*) [8]. We performed a similar ancillary analysis using full length sequences of Pax3/7 homologs, but recovered little phylogenetic structure within the gene tree (S3 Fig). Given that several chelicerate orders with unduplicated genomes retain more than one Pax3/7 homolog, we infer that a duplication of Pax3/7 genes likely predated the diversification of extant arthropods.

Taken together, our results suggest that *ey* and *toy* homologs were present in multiple chelicerate and pancrustacean exemplars, and by extension, across Arthropoda.

### Duplicates of *ey* and *toy* in arachnopulmonate genomes are attributable to whole genome duplication

A whole genome duplication in the common ancestor of Arachnopulmonata is evidenced by systemic paralogy across genetic elements, such as gene families, gene clusters (e.g., the Hox and Six genes) and microRNAs [37,39,45–47]. By contrast, previous surveys of Pax6 homologs across seven spider species have consistently reported both *ey* and *toy* as single-copy (in contrast to other RDGN members, such as *dachshund* [*dac*], *sine oculis/Six1* [*so*], *Optix/Six3*, and *orthodenticle* [*otd*]) [26,27,29]. We observed that four arachnopulmonates in our survey harbored two copies of *ey* (*Cordylochernes scorpioides* [Pseudoscorpiones], *Aphonopelma hentzi* [Araneae], *Phrynus marginemaculatus* [Amblypygi], and *Mastigoproctus giganteus* [Uropygi]). Two others retained a single copy of each (*Centruroides sculpturatus* [Scorpiones] and *Parasteatoda tepidariorum* [Araneae]). To test the hypothesis that the third Pax6 copy may reflect the process of whole genome duplication (albeit with the shuffling of gene order and loss of duplicated genes in one or the other chromosome), we assessed the identity of genes neighboring *ey* and *toy*.

We examined the identities of the 25 genes upstream of *toy* homologs and the 25 downstream of *ey* homologs in *P. opilio*, as well as four arachnopulmonates. We then circumscribed genes that were shared in this window in *P. opilio* and any arachnopulmonate. In the unduplicated genome of *P. opilio* (a non-arachnopulmonate arachnid), *ey* and *toy* occur on a single scaffold (ptg000048l_1) and are separated by a single gene (transmembrane 7 superfamily member 3), a homolog of which occurs upstream of a single *ey* homolog in *M. giganteus* (scaffold 12) (S4 Fig; S1 Table). Seven other genes occur in this 53-gene chromosomal window that are also found in at least one other arachnopulmonate scaffold containing a Pax6 homolog: *glucose dehydrogenase* (scaffold 8 of *M. giganteus*), *transposable element Tc1 transposase* (scaffold 7 of *A. hentzi*), *ETS homologous factor* (scaffold 7 of *A. hentzi*; scaffolds 8 and 12 of *M. giganteus*; scaffold 4 of *C. sculpturatus*; scaffold NC_092205.1 of *P. tepidariorum*), *DNA polymerase delta 1, catalytic subunit* (scaffold 12 of *M. giganteus*), *ankyrin repeat and SAM domain-containing protein 1A* (scaffolds 7 and 8 of *A. hentzi*; scaffolds 8 and 12 of *M. giganteus*; scaffold 4 of *C. sculpturatus*; scaffold NC_092205.1 of *P. tepidariorum*), *general transcription factor IIH subunit 1* (scaffold 12 of *M. giganteus*), and *buccalin* (scaffold 4 of *C. sculpturatus*). Comparing within the arachnopulmonates for similarly sized windows (51-55 genes; 25 genes upstream and downstream of Pax6 homologs), we discovered 17 other cases of homologs shared between at least two scaffolds.

We reasoned that the paucity of shared genes detected in the linkage group containing Pax6 homologs in the *P. opilio* genome (eight genes) could reflect gene shuffling and chromosomal reorganization over time, as *P. opilio* is the more distantly related to any arachnopulmonate than they are to each other. We therefore searched all five genomes for 19 genes that we identified as present in at least two species, and mapped these with respect to the Pax6 genes, regardless of the number of intervening genes. This broader search supported the inference of a shared linkage group. Fifteen out of 19 genes were present on the same scaffold as the Pax6 homologs in *P. opilio*, whereas the remaining three were retained on the same scaffold as a Pax6 homolog only in the arachnopulmonates (Fig 3; S2 Table). As measured by the number of genes retained on the same scaffold as a Pax6 homolog, the organization of the duplicated linkage group was better retained in *A. hentzi* and *M. giganteus* compared to *C. sculpturatus* and *P. tepidariorum*. While the two single-copy homologs of *ey* and *toy* both occurred on a single chromosome in the latter two species, we nevertheless found smaller fragments of the duplicated cluster on other scaffolds in *C. sculpturatus* and *P. tepidariorum*.

**Fig. 3.**
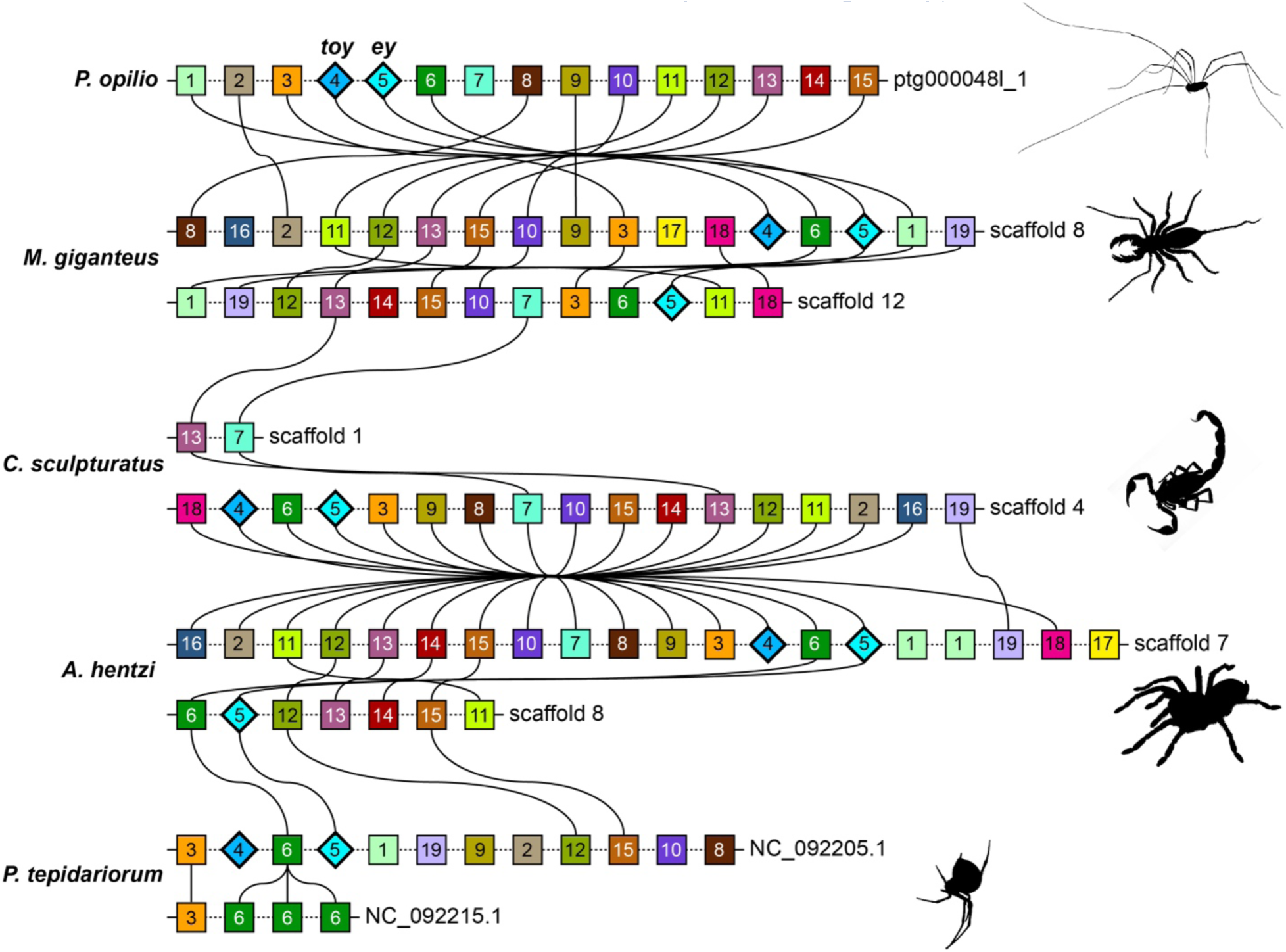
Conserved gene linkages support arachnopulmonate whole genome duplication as the mechanism underlying Pax6 paralogy. Chromosomal gene complements within 25 genes upstream and downstream of *Pax6* homologs. From top to bottom: *P. opilio* (Opiliones), *M. giganteus* (Uropygi), *C. sculpturatus* (Scorpiones), *A. hentzi* (Araneae, Mygalomorphae), and *P. tepidariorum* (Araneae, Araneomorphae). Identified genes are shared between at least two taxa. Ribbons represent homologs present in adjacent taxa. Gene identities: 1 – *zinc-cadmium resistance protein*; 2 – *BLOC-2 complex member HPS5/Hermansky-Pudlak syndrome 5 protein*; 3 – *ETS Homologous Factor*; 4 – *twin of eyeless*; 5 – *eyeless*; 6 – *ankyrin repeat/SAM domain-containing protein 1A*; 7 – *Buccalin*; 8 – *diuretic hormone class 2*; 9 – *syntaxin-12*; 10 – *prothoracicostatic peptide*; 11 – *troponin I*; 12 – *troponin T*; 13 – *sodium-dependent proline transporter*; 14 – *myosin kinase light chain, smooth muscle*; 15 – *ras associated domain-containing protein 10*; 16 – *fuzzy planar cell polarity protein*; 17 – *anaphase-promoting complex subunit 4*; 18 – *choline kinase alpha*; 19 – *serine/arginine-rich splicing factor 4*.

Taken together, the incidence of shared homologs in the same linkage group across arachnid orders, as well as the duplication of the linkage group in arachnopulmonates, is consistent with the scenario of a shared genome duplication in the common ancestor of Arachnopulmonata, in tandem with lineage-specific patterns of chromosome reorganization and gene loss.

### *ey* and *toy*, but not *sv*, are expressed in the head lobes during early *P. opilio* development

In contrast to spiders and horseshoe crabs, *ey* and *toy* are known to be expressed in the eyes of *P. opilio* throughout eye morphogenesis, with emphasis on lentigenic cells in later eye development [40]. However, expression of Pax genes in *P. opilio* is not known prior to stage 9. In the present study, HCR assays of earlier developmental stages detected *ey* and *toy* expression in the head lobes of stage 6 embryos (Fig 4A). The two homologs are co-expressed in two bilaterally symmetric domains in the protocerebral (first head) segment, with *toy* expression extending towards the ventral midline to form a continuous domain traversing the full segment (Fig 4A). *ey* is expressed in a smaller territory that co-expresses *toy*. At stage 7, *toy* is expressed discontinuously with respect to the ventral midline, with thin projections of *toy* expression still detected flanking the ventral midline (Fig 4B-C). As in stage 6, *ey* is co-expressed within a subdomain of *toy*. At stage 8, expression of both *ey* and *toy* broaden in the head lobes, expanding dorso-laterally. Additional bilateral regions of *ey* expression appear outside the head in each of the following body segments (Fig 4D). At this point, *toy* still retains a medial expression domain in the anterior most region of the head that lacks co-expression of *ey*.

**Fig. 4.**
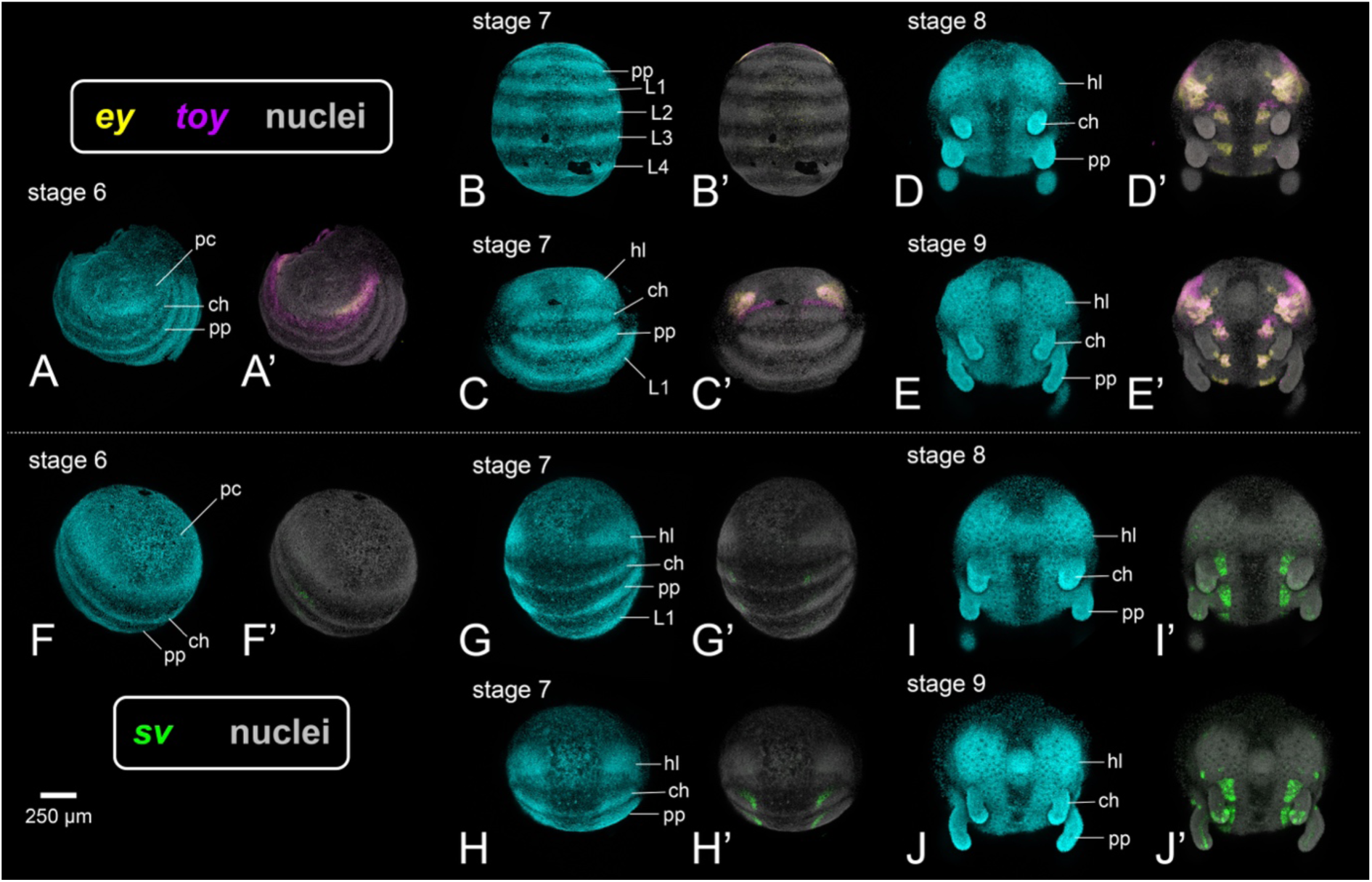
Expression of Pax homologs is consistent with a role for *ey* and *toy* (but not *sv*) in early head patterning. Top: HCR in situ hybridization for *Po-ey* (yellow) and *Po-toy* (magenta) in wild type embryos. (**A’-J’**) Expression patterns of targeted genes; (**A-J**) counterstaining of same embryos with Hoechst (cyan). (**A**) Stage 6. (**B**) Stage 7, view of ventral ectoderm. (**C**) Stage 7, view of head lobes. (**D**) Stage 8. (**E**) Stage 9. Bottom: HCR in situ hybridization for *Po-sv* (green). (**F**) Stage 6. (**G**) Stage 7, view of ventral ectoderm. (**H**) Stage 7, view of head lobes. (**I**) Stage 8. (**J**) Stage 9. Note the early expression of *Po-ey* and *Po-toy* in the head lobes by stage 7, whereas *Po-sv* remains restricted to ventral neurectoderm at the same stage. Abbreviations: pc, protocerebral segment; hl, head lobe; ch, chelicera; pp, pedipalp; L1, first walking leg.

Additionally, at stage 8, *ey* and *toy* expression in posterior segments appear as bilaterally symmetrical domains in the ventral ectoderm flanking the midline. At stage 9, expression of the two Pax6 homologs remains localized to the same regions as stage 8, but regions of heterogeneous expression strength result in a more complex expression pattern (Fig 4E).

Surveys of *sv* reveal a lack of expression in the ocular segment at stage 6 (Fig 4F). Instead, only bilaterally symmetrical bands of expression in the ventral ectoderm of posterior segments are observed. At stage 7, expression of *sv* intensifies as bilaterally symmetrical patches along each body segment but is absent from the protocerebral segment (Fig 4G, 4H). At stage 8, *sv* begins to be weakly expressed in small discontinuous territories in the lateral head lobes (Fig 4I).

Expression in the ventral ectoderm of posterior segments resolves into broad, bilaterally symmetrical domains. Additional domains are observed through the middle of each limb bud. At stage 9, *sv* expression in the limb buds resolves into clearly defined tracts of expression, corresponding to presumptive nerves (Fig 4J). Strong expression is observed in paired domains in the ventral ectoderm of each segment. In the head, *sv* is strongly expressed in the territory corresponding to the presumptive lateral eyes, as well as two domains at the anterior of the head lobes.

In summary, surveys of three Pax homologs’ expression suggest that (a) *ey* and *toy* exhibit conserved expression dynamics with respect to other arthropods, and (b) *sv* is not expressed in the protocerebrum until after the downstream RDGN genes *eya* and *so* have already commenced expression (stage 8; [40]).

### *ey* and *toy* are required for the development of all three eye pairs in *P. opilio*

No functional data exist for arthropod Pax genes outside of Pancrustacea. To redress this gap, we performed embryonic RNAi against *ey* and *toy* in *P. opilio*. To test the hypothesis that *sv* has gained a taxon-specific role in chelicerate eye patterning, we performed RNAi against the single-copy *sv* homolog of *P. opilio*. Phenotypes were scored in embryos at stage 17, by which point both well-formed eyes are present and the ozopores (repugnatorial gland pores) are first observable [42].

RNAi against *ey* elicited only wild type morphology (n = 154/225; 68.4%) or dead embryos (n = 71/225; 31.6%) (Fig 5A, 5B; S3 Table). Likewise, RNAi against *sv* elicited wild type eye development (n = 205/234; 87.6%) or dead embryos (n = 29/234; 12.4%). RNAi against *toy* resulted in a small number of embryos exhibiting subtle asymmetry of eye size, with one eye smaller than the other, but without evident defects in the morphology of the smaller eye (n = 14/271; 5.2%) (S3 Table). Penetrance of knockdown was measured using RNA sequencing, which supported statistically significant reduction of target gene expression level in all single-gene experiments (Fig. 5C; S5 Fig; S4 Table). Knockdown of *ey*, *toy*, or *sv* resulted in significant reduction of expression in all three Pax genes, except for *ey* RNAi, which had no effect on the expression of *Pax2* relative to control experiments (S5 Fig). Knockdown of *ey*, *toy*, or *sv* also generally resulted in reduction of other Pax gene family members and downstream RDGN genes, with the notable exception that RNAi against *toy* resulted in significant upregulation of *Optix* expression (S5 Fig).

**Fig. 5.**
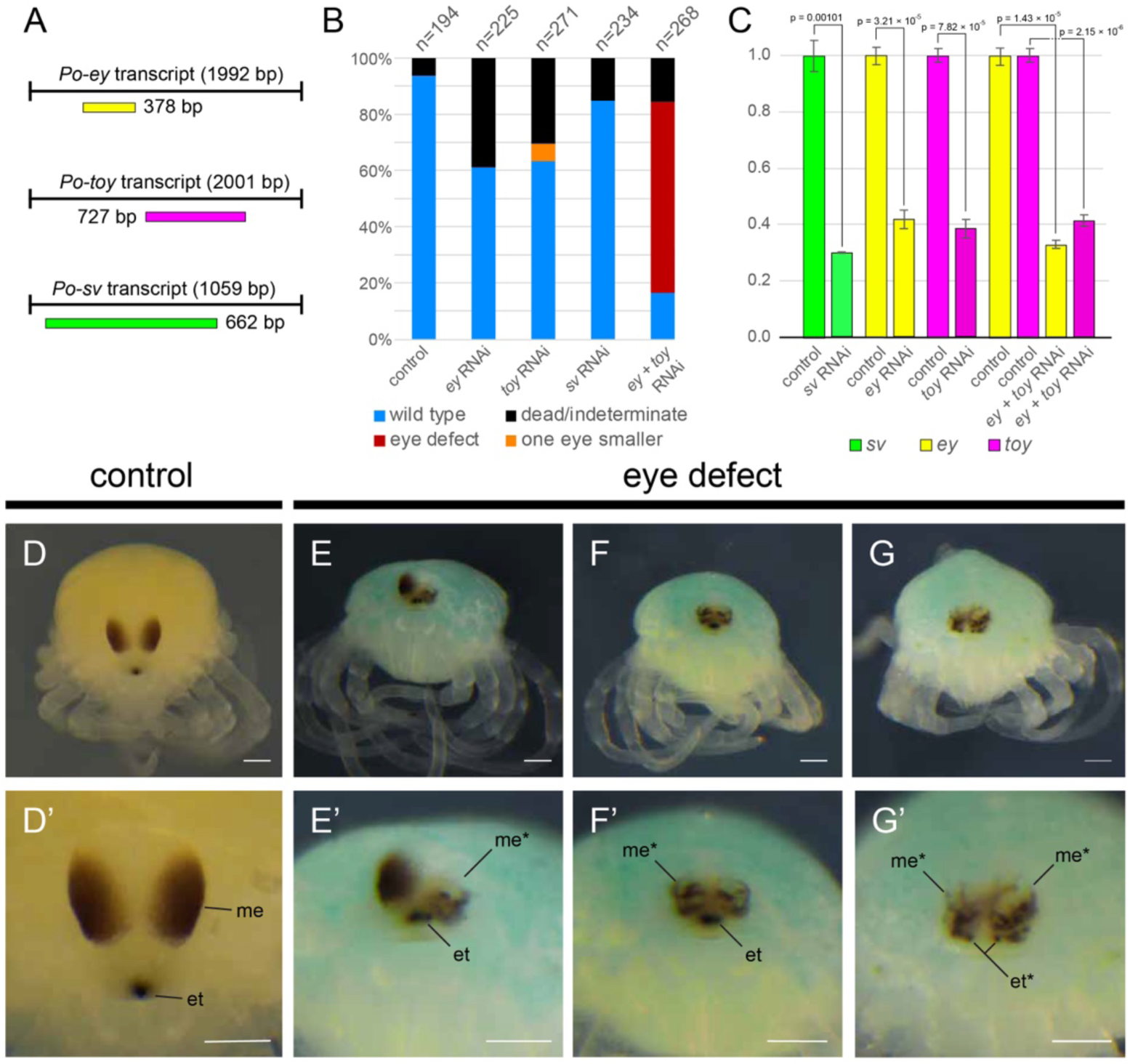
Double knockdown of *Po-ey* and *Po-toy* disrupts median eye patterning in *P. opilio*. (**A**) Locations and lengths of cDNA fragments amplified from *ey*, *toy*, and *sv* transcripts for synthesis of dsRNA in RNAi experiments. (**B**) Distribution of phenotypic outcomes following RNAi experiments. (**C**) Relative expression of *sv*, *ey*, and *toy* transcripts for control and RNAi embryos. (**D**) Brightfield image of stage 17 wildtype *P. opilio* embryo showing normal eye development. (**D’**) Same embryo as in (**D**) with magnification of median eyes. (**E-G**) Bright-field images of spectrum of eye defects in double *Po-ey*+*Po-toy* RNAi embryos. (**E**) RNAi embryo with one small median eye, one defective median eye, and normal egg tooth. (**F**) RNAi embryo with two defective median eyes and normal egg tooth. (**G**) RNAi embryo with two defective median eyes and defective egg tooth without fusion in the midline (**E’-G’**) Same embryos as in (**E-G**), with magnification of median eyes. Note the mosaic phenotype in (**E**) affecting only one of the two median eyes. Abbreviations: et, egg tooth; et*, egg tooth with developmental defect; me, median eye; me* median eye with developmental defect. Scale bars: 100 µm.

It has previously been shown that *ey* and *toy* act synergistically in the patterning of the fruit fly and beetle compound eye; joint knockdown of both Pax6 genes induces a stronger effect than single-gene knockdown, which is inferred to result from functional redundancy between the two homologs [2,16]. We therefore performed double-knockdown of *ey* and *toy*, which elicited a spectrum of defective eye phenotypes. RNAi embryos exhibited eyes that were smaller, heterogeneously pigmented, and missing components of the cuticular lens (n = 148/268; 88.1%) (Fig 5E-5G; S3 Table). RNAi embryos with the most severe eye defects exhibited a defective egg tooth that did not fuse in the midline, a smaller ocularium (the eye mound that bears the median eyes), and anophthalmia; such embryos did not survive to later stages of development and were fixed for HCR assays (below).

Our recent discovery of vestigial eye pairs in *P. opilio* facilitates an investigation of Pax gene function with respect to the two visual systems of chelicerates. We surveyed the expression of three opsins that are differentially expressed in the three eye pairs as readouts of proper eye morphogenesis: *peropsin* (median eyes), *Rh1* (vestigial median and vestigial lateral eyes), and *Rh3* (vestigial median and vestigial lateral eyes) [40]. Expression assays showed a spectrum of phenotypes, with opsin expression only in vestigial lateral eyes (n = 8/12) or lost in all three eye pairs (n = 4/12), with the latter associated with embryos that bore drastically reduced ocularial height (Fig 6). Consistent with the high degree of mosaicism in eye morphology of RNA embryos at stage 17, we observed a spectrum of mosaicism in HCR surveys, with one side visibly more strongly affected than the other (S6 Fig).

**Fig. 6.**
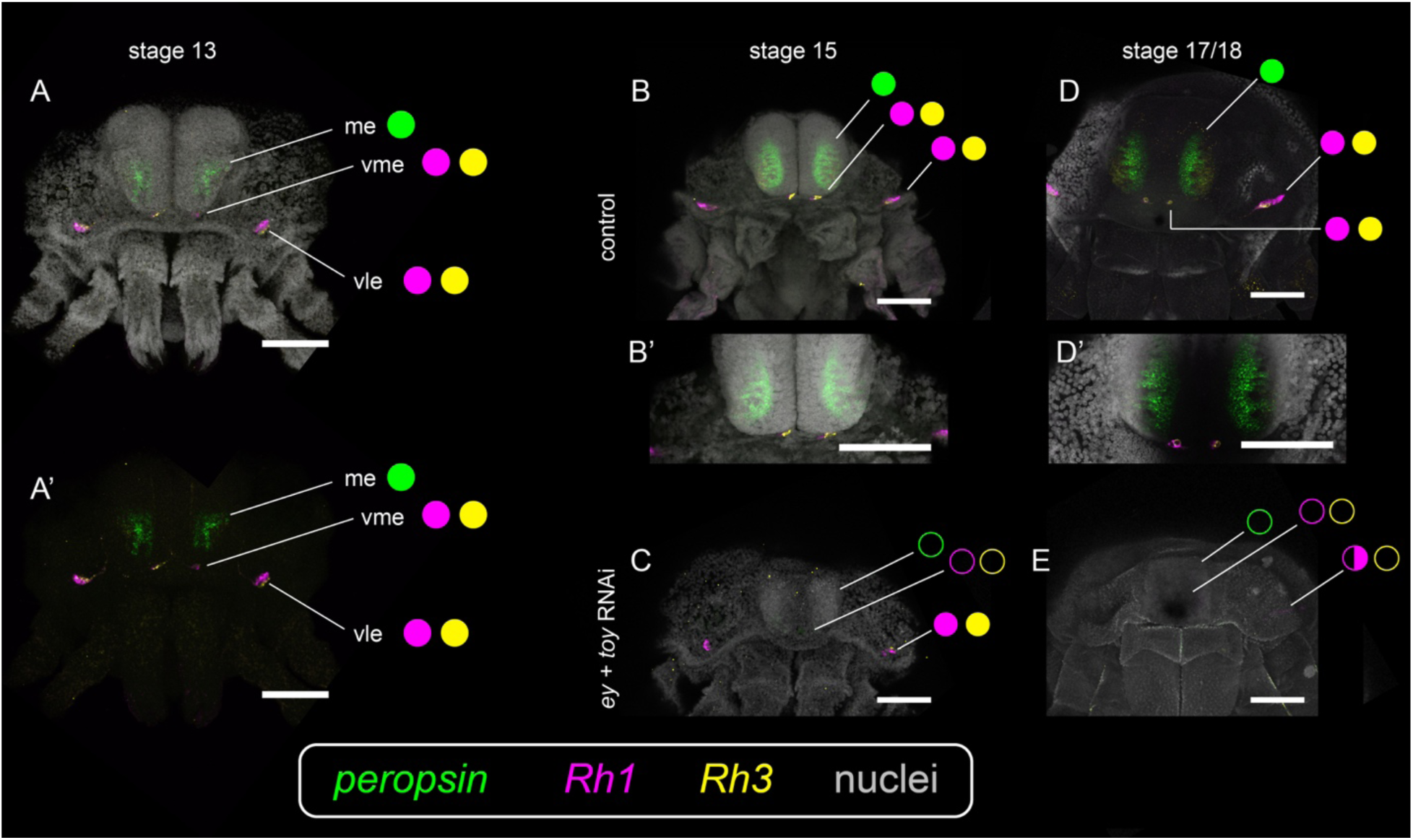
RNAi against *Po-ey*+*Po-toy* disrupts development of both median and lateral visual systems. (**A-E**) Hoechst counterstaining (gray) and multiplexed expression of *Po-peropsin* (green), *Rh1* (magenta), and *Rh3* (yellow) in *P. opilio* embryos. (**A**) Wild type embryo at stage 13. (**A’**) Same embryo as in (**A**) with isolated opsin expression. Note that the expression of *Po-peropsin* is restricted to the median eyes, whereas *Rh1* and *Rh3* are restricted to the vestigial median and vestigial lateral eyes. (**B**) Stage 15 negative control embryo. (**B’**) Same embryo as in (**B**) with magnification of the median eyes. (**C**) Stage 15 *Po-ey+Po-toy* RNAi embryo. (**D**) Stage 17 or 18 negative control embryo. (**D’**) Same embryo as in (**D**) with magnification of median eyes. (**E**) Stage 17 or 18 *Po-ey+Po-toy* RNAi embryo. Note the loss of *Po-peropsin* expression in the median eyes, and loss of *r-opsin* and *Rh3* in the vestigial median eyes in RNAi embryos. Filled circles represent wildtype expression of opsins in each eye pair. Empty circles represent complete absence of opsin expression. Half-filled circles represent mosaic phenotypes (retention of wild type expression on only one side). Abbreviations: ME, median eyes; VME, vestigial median eyes; VLE, vestigial lateral eyes. Scale bars: 100 µm.

To address the possibility of differential roles for *ey*, *toy*, or *sv* in subdividing the patterning of different eye types, we performed the same HCR assays for single-gene knockdown experiments. However, all embryos assayed in these experiments exhibited wild type expression of all three opsins (n = 65).

Beyond eye patterning defects, the insect Pax6 loss-of-function phenotypic spectrum includes defects of the head that include microcephaly, as Pax6 plays an additional role in head and brain development. For example, a maternal RNAi experiment in the flour beetle *Tribolium castaneum* showed that reduction of *dachshund* expression in the compound eye in *ey+toy* RNAi embryos is the result of a deletion of that territory of the developing head lobe [7]. CRISPR experiments in a honeybee have also shown that *toy* knockout results in eye defects and microcephaly [48]. To assess the possibility that loss of opsin expression in the vestigial median eyes in *P. opilio ey*+*toy* RNAi phenotypes could have resulted from secondary effects of head deletion, we assayed RNAi embryos for the RDGN members *dac*, *so*, and *eya*. *ey+toy* RNAi embryos exhibited diminution of *eya* in the head lobes, consistent with a known anophthalmia phenotype previously described for *eya* RNAi in *P. opilio* (Fig 7). These embryos also exhibited diminution of *so* and *dac* which is consistent with known roles of these genes in patterning both visual systems of a spider and the lateral visual system in *P. opilio*, respectively [31,40]. HCR is challenging when interpreting degree of knockdown of transcription factors; we therefore examined quantitative expression of these genes, using an RNA-Seq dataset. Consistent with the imaging data, normalized counts of these RDGN members showed significant knockdowns in all three genes (independent two-sample, two-tailed t-tests; p << 0.05). Nevertheless, we detected no qualitative differences in the expression patterns of these three genes, disfavoring the interpretation of head deletions in *ey*+*toy* RNAi embryos (Fig 7). This is consistent with the implementation of our embryonic RNAi experimental design; the stages injected with dsRNA in this study have already formed the head lobes (stage 6 of *P. opilio*; [42]), and thus *ey+toy* RNAi should not affect earlier head-patterning functions, in contrast to maternal RNAi experiments in *T. castaneum*.

**Fig. 7.**
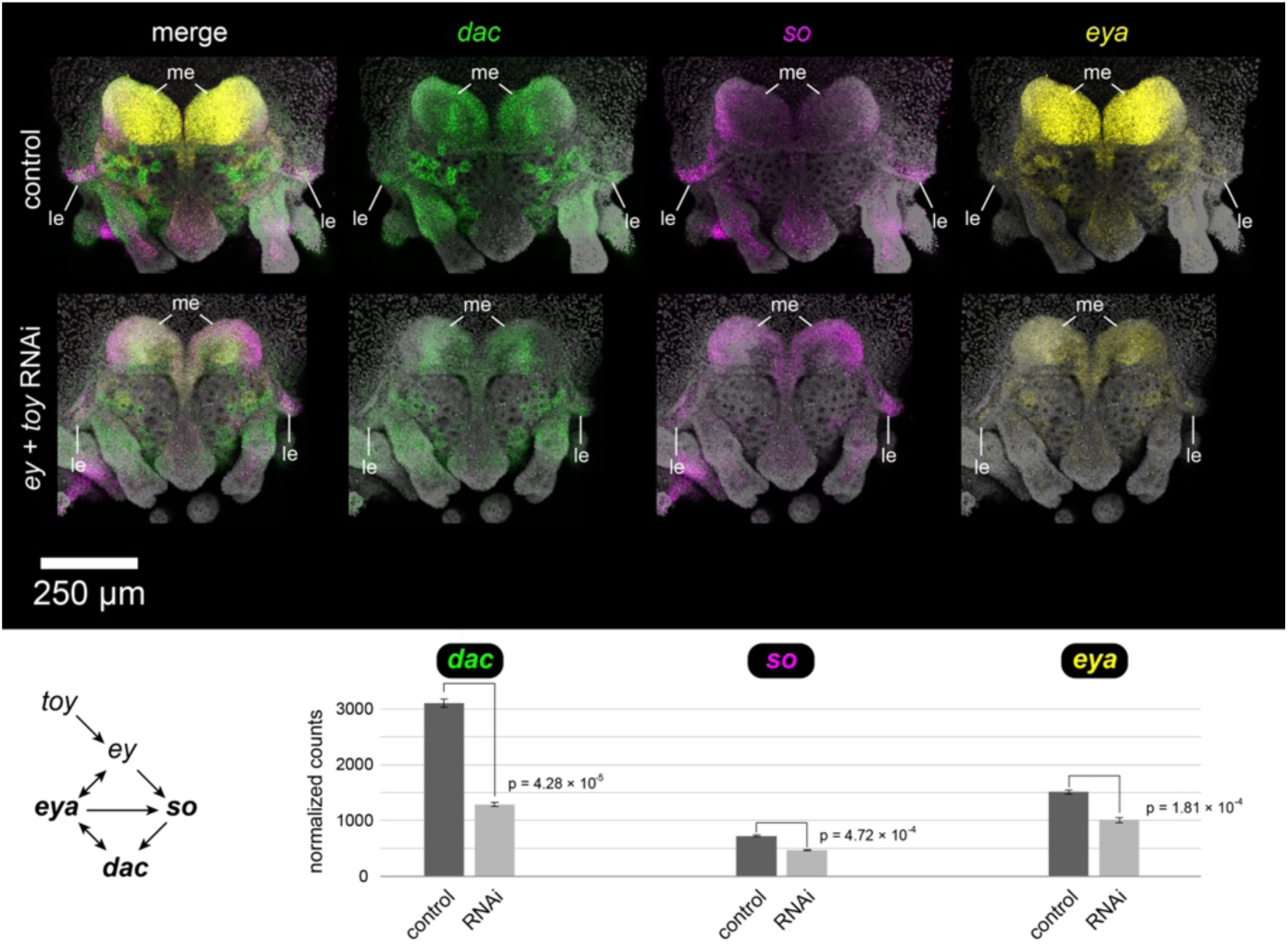
Embryonic *Po-ey*+*Po-toy* RNAi reduces target gene expression, but does not result in deletions of head territories. HCR assays for *dac* (green), *so* (magenta), and *eya* (yellow). Gray: Hoechst counterstain. Top row: Negative control embryo 72 hr after injection. Middle row: *Po-ey*+*Po-toy* RNAi embryo 72 h after injection. Note the diminution of *dac*, *so*, and *eya* expression, though the morphology of the embryo does not support the deletions of head regions, with emphasis on the lateral eye territory (le). Bottom row: Gene regulatory network showing present understanding of RDGN regulatory interactions, after [7]; and normalized read counts of surveyed genes 48 h after injection. *p* values correspond to independent two-sample, two-tailed t-tests. Abbreviations as in Fig 4.

Taken together, our results are consistent with the understanding that *ey* and *toy* act synergistically to pattern both visual systems of an arachnid, closely paralleling the results of insect experiments [2]. By contrast, we found no role for *sv* in patterning any of the eyes, consistent with its later onset of expression in the head in stages after the onset of *so* and *eya* expression.

## Discussion

### A conserved role for Pax6 homologs in the patterning of the two arthropod visual systems

Despite long-standing interest in the evolution of the arthropod visual system, the developmental genetic literature harbors a prominent dearth of functional data from phylogenetically significant groups, such as chelicerates, myriapods, and Onychophora (the sister group to Arthropoda). Part of this gap is attributable to the complete absence of functional tools in exemplars of Myriapoda and Onychophora, but this limitation does not apply to chelicerates. RNA interference has been available as a tool for the study of developmental genetics in spider models for 25 years, though its efficiency is variable across studies [30,32,49–51]. For example, our previous RNAi study on *so-1* in the spider *P. tepidariorum* was able to elicit a spectrum of anophthalmia phenotypes, albeit in only 9.5% of embryos [31]. Despite this seemingly promising beginning, we and others have also previously performed RNAi against every RDGN gene duplicate in *P. tepidariorum* as single-gene knockdowns; none of these experiments yielded eye defects [31,52]. As there are no functional data corresponding to any other spider eye-patterning genes beyond *so-1*, most of the present understanding of spider eye-patterning (e.g., RDGN ohnolog dynamics; Hedgehog and Wnt signaling) is based on interpretation of gene expression patterns in tandem with inferences of function derived from the fruit fly literature [29,53,54]. Advanced functional tools (e.g., maternal CRISPR-mediated knockout and knock-in) do exist for some chelicerate models, such as the mite *Tetranychus urticae* and the tick *Ixodes scapularis*, but these models either lack eyes (*I. scapularis*) or ongoing work on these systems has mostly been directed toward advancing study of biological control in support of agricultural and biomedical interests [55–57]. The recent establishment of a CRISPR technique in the spider *P. tepidariorum* is similarly not promising without further development, due to intractably low penetrance and the inability of the embryos to survive to hatching in the injection medium, precluding the establishment of transgenic lines or full characterization of structures that develop post-embryonically [58], including the eyes [31].

This gap in our understanding of arachnid eye development has been bridged by the daddy-longlegs *P. opilio*, both due to the efficiency of RNAi in this species as well as the recent discovery of a plesiomorphic visual architecture featuring two sets of vestigial eyes in Opiliones. We therefore brought this system to bear upon the question of Pax gene function in Chelicerata.

Both the expression pattern and the lack of an eye-patterning function of *sv* disfavor the hypothesis that Pax2 has superseded Pax6 function in eye fate specification (either median or lateral). This is consistent with the known function of *sv* in *D. melanogaster*, wherein *sv* is necessary for the differentiation of cone and primary pigment cells and regulating crystallin production [59]. Mutants for *sv* exhibit defects in ommatidial cell development, which include cell fate transformations and a disruption in the formation of proper cell structures. These known functions accord with the onset and expression patterns of Pax2 homologs in embryonic daddy-longlegs, spiders, and scorpions [36,40]. Our previous surveys have also shown that the subdivision of expression patterns of Pax2 ohnologs in spiders is taxon-specific; in a scorpion (two *sv* ohnologs) and a harvestman (single-copy *sv*), *sv* homologs are expressed in both the median and lateral eye primordia. These expression surveys tentatively support a series of derived events in some lineage subtending spiders (e.g., crown-group Araneae; Tetrapulmonata), where that lineage lost *Pax2.1* expression in the median eyes and *Pax2.2* expression in the median and lateral eye [41]. These spider-specific expression dynamics are therefore intriguing, but their significance remains known. The availability of new genomic and developmental resources for exemplars of Pedipalpi (the sister group of Araneae) at least offers a potential means to pinpoint this evolutionary sequence of expression domain changes across the arachnopulmonates [60].

Like *sv* RNAi, single-gene knockdown experiments against *ey* and *toy* did not elicit an eye defect phenotype (though a minority of *toy* RNAi embryos exhibited mild asymmetry of eye size). By contrast, double-knockdown of *ey* and *toy* elicited a consistent spectrum of eye defects with >85% efficiency. In these double RNAi experiments, expression assays of differentially expressed opsins corroborated defects to the vestigial median and vestigial lateral eyes as well, suggesting that *ey* and *toy* act synergistically to pattern all three eye pairs of *P. opilio*. The significance of this result is that it closely parallels Pax6 knockdown experiments in insects, with single-knockdown RNAi experiments eliciting limited phenotypic spectra and double knockdown of *ey*+*toy* eliciting a range of microphthalmic and anophthalmic phenotypes with higher penetrance [2]. The requirement that both *ey* and *toy* are silenced to obtain this phenotypic spectrum is consistent with the understanding that these two Pax6 homologs can reciprocally rescue one another’s loss of function and that they compete for binding sites of target genes [16]. In addition, the evolutionary significance of these results is that they support an ancestral requirement of *ey* and *toy* for patterning both median and lateral visual systems of insects and arachnids, and by extension (given the phylogenetic placement of chelicerates as the sister group of the remaining arthropods), across the phylum.

Recent advances in the understanding of Pax6 function in the fruit fly have leveraged the availability of functional tools to demonstrate how precise expression levels of *ey* and *toy* establish ocellar and compound eye fates in *D. melanogaster*. A remarkable component of these experiments achieved the homeotic transformation of the ocelli to compound eyes through overdriving expression of *eya*, a downstream target of Pax6, in the anterior head via a GAL4 driver [16]. The significance of this study is that it connects the dissimilar morphologies of ocelli and facetted eyes as part of an evolutionary transformation series, underpinned by a shared developmental program, despite hundreds of millions of years since the separation of the two visual systems. It has been suggested that this mechanism may be conserved across arthropods as well, a possibility that could be tested by altering levels of Pax6 homolog expression in the median versus lateral eyes of spiders [16]. Due to the unavailability of advanced functional tools in spider and daddy-longlegs models, overdriving expression in specific tissues is not feasible in these models, which are presently limited to gene knockdown approaches. Nevertheless, a potential route to testing conservation of eye-patterning mechanisms with available tools may be to trial RNAi against *groucho* (*gro*), a broadly expressed transcriptional co-repressor that has been shown to mediate the choice between ocellar and compound eye fate in *D. melanogaster* [16]. Specifically, *gro* mutants and GAL4-driven *gro* RNAi lines exhibit hyperproliferation of ocellar field or transformation of the anterior head epidermis into ocelli. This effect is additive; GAL4-driven overexpression of *eya* in the anterior head in tandem with *gro* RNAi results in both homeotic ocelli-to-compound eye transformations, as well as homeosis of the head epidermis in between ectopic compound eyes into ocelli [16]. Future investigation of eye-patterning in *P. opilio* should therefore trial *gro* RNAi, with the prediction that abrogation of *gro* expression levels should incur hypertrophy of the median eye fields.

### Genome architecture of the linkage group containing *ey* and *toy* supports an arachnopulmonate WGD event

A recent survey of RDGN genes in spiders reported single-copy homologs of both *ey* and *toy* in all surveyed taxa, which included the theraphosid *Acanthoscurria geniculata* [29]. We were therefore surprised by the occurrence of three Pax6 copies in the genomes of the theraphosid *A. hentzi*, as well as two other arachnopulmonate species (*M. giganteus* and *C. scorpioides*). A possible explanation for this third copy is that it represents the signature of the whole genome duplication (WGD) event shared in the common ancestor of Arachnopulmonata; the requirement of precise expression levels of Pax6 during eye fate specification may have selected against the retention of duplicate copies of *ey* and *toy*, resulting in widespread purging of their duplicates across arachnopulmonate genomes. However, one study has refuted the incidence of a WGD in arachnids altogether [61]. The basis for this conclusion was that (a) distributions of interparalog distances (Ks plots) in selected arachnopulmonates were not consistent with WGD, (b) only a subset of genes such as the Hox genes exhibited systemic paralogy, (c) self-synteny plots did not support patterns of duplication within arachnopulmonate genomes, and (d) arachnopulmonate genomes did not exhibit syntenic patterns supporting WGD, as inferred from blocks of contiguous genes with conserved gene order.

As discussed in our previous analysis of the *M. giganteus* genome [62], (a) interparalog distances inferred using simplistic nucleotide substitution model corrections are deeply sensitive to model choice and lose power to detect ancient WGD events as a function of WGD event age [63]; (b) systemic paralogy in arachnopulmonate genomes is widely known to extend well beyond the Hox genes and is substantiated by analyses of various gene families, gene expression patterns, and microRNAs [39,45–47,64,65]; (c) self-synteny has been reliably recovered in multiple arachnopulmonate (and particularly spider) genomes [62,66,67]; and (d) a low discovery rate of syntenic patterns is closely tied to the use of conserved gene order as a defining criterion [68]. For example, a syntenic block may be contextually defined in a given study as a minimum of three adjacent genes that retain a conserved gene order (e.g., A-B-C; C-D-E). The problem with this definition is that even a single rearrangement within a small block of genes (e.g., A-B-C-D-E to A-B-E-D-C) will break the required minimum of three genes retaining the same gene order. Such a narrow definition of synteny for defining linkage groups works well for recent WGD events, but rapidly accrues a high false negative rate as the age of the WGD event increases. This is due to the accumulation of gene shuffling and genomic rearrangements over evolutionary time, particularly among small-bodied and rapidly evolving arthropod groups [47,69]. Indeed, a return to the classic definition of synteny—gene linkages without consideration for gene order—has dramatically augmented the ability to detect ancient duplicated chromosomes in the metazoan (and especially vertebrate) tree of life [68,70]. The application of this concept of synteny to Arachnopulmonata and Xiphosura has accelerated the identification of duplicated ancestral chromosomes, though there is substantial variability in the cohesion of syntenic patterns across arachnopulmonate taxa [47,62]. Specifically, lineages that exhibit longer generation times and lower evolutionary rates (e.g., mygalomorph spiders; Uropygi) tend to retain stronger signatures of WGD than lineages with short generation times and high evolutionary rates (e.g., small-bodied araneomorph spiders) [47].

Our examination of genome architecture in the context of the third Pax6 gene copy in *A. hentzi* and *M. giganteus* supported the inference of paralogy via WGD. When the criterion of gene order is imposed on arachnopulmonates, little evidence is available for syntenic blocks containing Pax6 homologs. When the criterion for gene order is relaxed, ancestral gene linkages harboring the Pax6 homologs are evident, as are conserved arrangements of genes within those linkage groups, albeit with additional genes interspersed between identified ohnologs. These results support the interpretation that various embryonic-patterning genes and gene families may not faithfully retain duplicated copies stemming from the arachnopulmonate WGD (contra well-studied cases such as the homeobox and Wnt gene families; [46,65,71]), possibly owing to different selection pressures acting upon different developmental processes. Moreover, the identification of numerous putative ohnologs that are unrelated to embryonic patterning suggests that the true scale of paralogy in arachnopulmonate genomes has not yet been fully explored. Ongoing establishment of new genomic resources for phylogenetically significant groups, in tandem with exploration of ancestral gene linkages specific to Chelicerata, are anticipated to improve the understanding of how ohnologs are retained or lost in lineage-specific patterns.

A corollary of these observations is that the persistence of duplicated genes in arachnopulmonates may not reflect adaptive processes such as subfunctionalization or neofunctionalization, which are frequently invoked as explanatory vehicles for the diversification of arachnopulmonates. Specifically, divergence of expression patterns in arachnopulmonate ohnologs is often associated with divergence of function, morphological diversification, and/or the emergence of novel traits, though functional support for these associations are frequently lacking [26,36,72,73]. A valid and competing hypothesis is that subsets of duplicated genes retained after WGD may simply be redundant, as exemplified by recent cases of posterior patterning in spiders [58,74], or even in the process of gradual loss, a process that has received recent theoretical attention [75]. Functional redundancy between persisting gene copies may also explain the recalcitrance of spider RDGN duplicates to RNAi. One potentially unexplored model for such investigations may be the horseshoe crabs, whose genomes retain a higher proportion of ohnologs, likely due to much younger genome duplications [76–79]. Investment in developing functional tools for Xiphosura may yield compelling insights about the incidence and persistence of functional redundancy between ohnologs, particularly in the context of *ey* and *toy* paralog dynamics during median versus compound eye development in this chelicerate order.

## Conclusion

The role of Pax6 homologs in Chelicerata has long been a topic of interest, given expression data periodically generated for spiders and horseshoe crabs, but these discussions have been hampered by a lack of functional data. We were able to bridge this gap by generating a loss-of-function phenotype via double knockdown of *ey* and *toy*. This dataset, the first of its kind for Chelicerata, supports the inference that Pax6 genes play an evolutionarily conserved role in patterning both visual systems of an arachnid and insects, and by extension, across the arthropods. Next steps in deciphering the evolutionary history of arthropod eye evolution should prioritize testing the hypothesis that duplication and functional subdivision of *ey* and *toy* drove the origin of lateral and median visual systems in the common ancestor of Arthropoda.

## Materials and Methods

### Gene tree analysis and orthology inference

To survey Pax homologs across Chelicerata, 13 chelicerate genomes or developmental transcriptomes were selected for comparative analysis, as well as six outgroup datasets. The 13 chelicerate genomes surveyed comprised *Parasteatoda tepidariorum* (Araneae), *Aphonopelma hentzi* (Araneae), *Phrynus marginemaculatus* (Amblypygi), *Mastigoproctus giganteus* (Uropygi), *Centruroides sculpturatus* (Scorpiones), *Cordylochernes scorpioides* (Pseudoscorpiones), *Carcinoscorpius rotundicauda* (Xiphosura), *Gluvia dorsalis* (Solifugae), *Titanopuga salinarum* (Solifugae), *Tetranychus urticae* (Acariformes), *Ixodes scapularis* (Parasitiformes), *Phalangium opilio* (Opiliones), *Pycnogonum litorale* (Pycnogonida). The outgroup genomes comprised *Drosophila melanogaster* and *Daphnia pulex* (Mandibulata), *Euperipatoides kanangrensis* and *Epiperipatus broadwayi* (Onychophora), and *Rammazottius varieornatus* and *Paramacrobiotus metropolitanus* (Tardigrada).

BLASTp or tBLASTn searches were performed on these 19 datasets using the PRD domain of the ten *D. melanogaster* Pax genes as peptide queries. All hits with e-values < 10^-10^ were retained. All putative orthologs were verified using reciprocal BLAST searches and gene tree analysis. Multiple sequence alignment was conducted using CLUSTAL Omega [80].

Phylogenetic reconstruction consisted of maximum likelihood analysis with IQ-TREE v3 [81], with automated model selection (-m MFP; chosen model: LG +G4) and 1,000 ultrafast bootstrap resampling replicates. Three families of analyses were performed. First, a gene tree was inferred for all Pax genes using only the PRD domain in the multiple sequence alignment. Next, to decipher relationships between *ey* and *toy*, a gene tree was inferred using full length sequences of all Pax6 homologs without trimming. Third, to decipher relationships between Pax3/7 homologs, a separate gene tree was inferred using full length sequences of all Pax3/7 homologs without trimming.

### Analyses of synteny

To test for patterns of conserved gene linkages, scaffolds containing *ey*, *toy*, or both were annotated to determine the identity of neighboring genes in the genome assemblies of *P. opilio*, *A. hentzi*, *P. tepidariorum*, *M. giganteus*, and *C. sculpturatus*. The *P. opilio* genome was used as a starting point, due to its unduplicated condition. On the scaffold containing *ey* and *toy*, the 25 genes upstream of *toy* and the 25 genes downstream of *ey*, as well as a single gene in between *ey* and *toy*, were all identified using best reciprocal BLASTp hit. The same approach was repeated for the remaining four species (25 genes upstream and downstream of *ey* and *toy* homologs). Genes with identical BLAST hits in any two scaffolds were identified as homologous cliques.

To assess the possibility of more extensive shuffling of genes, BLASTp searches were then performed for each clique, for all five arachnid species. Locations of any BLAST hits matching scaffolds containing *ey* or *toy* were retained. The relative order of genes was established using GFF or GTF files for the corresponding genome.

### Collection of embryos and gene expression assays

Embryos of *P. opilio* were collected using coconut fiber dishes, as previously described [42]. Stage 6-10 embryos were fixed and assayed for hybridization chain reaction. Staging of embryos and protocols for fixation follows a recent staging system [42].

Orthology inference for *Po-ey*, *Po-toy*, *Po-sv*, *Po-eya*, *Po-so*, and *Po-dachshund*, design of hybridization chain reaction (HCR) probes, and protocols for fluorescent detection of gene expression were previously reported in our prior work [40]. The same HCR probes were redeployed to survey earlier stages of *P. opilio* development.

### RNA interference

Primers were designed for fragments of the coding regions of *Po-ey*, *Po-toy*, and *Po-sv* using Primer3 (S5 Table). These fragments were amplified using standard PCR protocols and cloned using a TOPO TA Cloning Kit using One Shot Top10 chemically competent *Escherichia coli* (ThermoFisher) following the manufacturer’s protocol. Amplicon identity was verified via Sanger sequencing with M13 universal primers. Double-stranded RNA (dsRNA) was synthesized following the manufacturer’s protocol using a MEGAscript T7 kit (Ambion/Life Technologies) from PCR templates. The quality of dsRNA was assessed and concentrations adjusted using a NanoDrop ONE to 3.9-4.1 µg/µL. dsRNA was mixed with vital dyes for visualization of injections. For double knockdown, dsRNA for each gene was adjusted to a concentration of 6.0 µg/µL and combined in equal volume. As negative controls, we used dsRNA for an exogenous fragment of vector plasmid sequence amplified using M13 universal primers. Microinjection under halocarbon-700 oil (Sigma-Aldrich) was performed as previously described at the peri-vitelline space stage. Subsets of injected embryos were fixed for HCR; the remainder was developed at 26°C until stage 17-19 for scoring eye phenotypes.

### Validation and differential gene expression analyses

Sets of 30-35 embryos were injected with dsRNA for *sv*, *ey*, *toy*, *ey + toy*, or an exogenous vector sequence. Embryos were fixed 48 h after injection using TriZOL trireagent (ThermoFisher). First-strand cDNA was prepared using oligo(dT) primers and Super-Script III reverse trancriptase (ThermoFisher).

qPCR was trialed using PowerUp SYBR Green Master Mix (Life Technologies) on a StepOne Plus Real-Time PCR instrument (Applied Biosystems) following the manufacturer’s protocol and using primers for elongation factor 1α as the endogenous control (qPCR primers in Table S2). However, low C_t_ values despite high input cDNA (>35 cycles for target genes of control embryos in pilot assays) precluded this strategy for validation of the knockdown.

As an alternative, RNA sequencing was performed using total RNA extractions of sets of 30-35 embryos at 48 hours after injection. Library preparation was performed using an Illumina TruSeq kit, following manufacturer’s protocols. Sequencing. Stranded mRNA sequencing was performed on an Illumina NovaSeq with 2x150 bp paired end sequencing.

*Phalangium opilio* genome assembly version 2 (GenBank accession no. GCA_054372755.1) was converted into a reference index using STAR v.2.7.11 [82], followed by sequential mapping of each read file to the genome. Analysis of differential gene expression was performed using DESeq2 v. 1.48.2 [83]. DESeq2 was run with cutoff values for identification of differentially expressed genes set to a p-value of < 0.05 and log-fold change > 1.

### Imaging

Brightfield microscopy was performed using a Nikon SMZ fluorescence stereomicroscope mounted with a DSFi2 digital color camera and driven by Nikon Elements software. Images of *P. opilio* appendages were captured at varying focal planes and compiled into focused stacks with inbuilt software tools. Confocal laser scanning microscopy was performed using a Zeiss LSM 980 microscope driven by Zen software. Maximum intensity projections were assembled using inbuilt software tools in Fiji [84].

## Acknowledgements

This work was supported by National Science Foundation grant no. IOS-2016141 to PPS. EML was supported by a Hilldale undergraduate research scholarship award. Imaging infrastructure was provided by the Newcomb Imaging Center, Department of Botany, University of Wisconsin-Madison. Discussions with Guilherme Gainett and Markus Friedrich refined the ideas presented in this paper.

## Supplementary figure and table legends

**Table S1.** BLASTp hits for genes adjacent to *ey* and *toy* on scaffolds of representative arachnids.

**Table S2.** Locations of targeted genes adjacent to *ey* and *toy* on scaffolds of representative arachnids.

**Table S3.** Phenotypic distribution from RNAi experiments.

**Table S4.** Normalized counts and relative expression of Pax and RDGN genes.

**Table S5.** Primer sequences used in this study.

**Fig. S1.** Maximum likelihood tree topology inferred from multiple sequence alignment of the PRD domain. Numbers on nodes indicate ultrafast bootstrap resampling frequencies.

**Fig. S2.** Maximum likelihood tree topology of full length Pax6 sequences without trimming. Numbers on nodes indicate ultrafast bootstrap resampling frequencies.

**Fig. S3.** Maximum likelihood tree topology of full length Pax3/7 sequences without trimming. Numbers on nodes indicate ultrafast bootstrap resampling frequencies.

**Fig. S4.** Gene identities 25 genes upstream and downstream of *Pax6* homologs (rhombus icons). Black squares indicate loci with BLAST hits that do not share homologs in other scaffolds. Gray squares indicate loci with low-confidence or no BLAST hits (e.g., “hypothetical protein”; “unidentified protein”).

**Fig. S5. (A)** Principal components analysis clustering of RNA sequencing samples. **(B)** Hierarchical clustering of RNA sequencing samples for top 200 differentially expressed genes. **(C)** Relative gene expression of Pax family members and downstream RDGN genes in RNA sequencing experiments. Asterisks indicate significance of two-tailed t-tests (p < 0.05).

**Fig. S6.** RNAi against both *Po-ey* and *Po-toy* yields on-target reductions in expression. Stage 9 *Po-ey*+*Po-toy* RNAi embryo with Hoechst nuclear counterstaining (gray) and multiplexed expression of *Po-ey* (green) and *Po-toy* (magenta). Note the differences in expression strength for both *Po-ey* and *Po-toy* between the two halves of the embryo, which is consistent with mosaicism of knockdown.

