## Supplementary material for "Pax6 homologs are required for patterning both visual systems of the daddy-longlegs *Phalangium opilio*": Table S3

**Table S3. Distribution of phenotypes from RNAi experiments.**

|  | wild type | one eye asymmetrically sized | eye defect | dead/indeterminate | Total |
| --- | --- | --- | --- | --- | --- |
| control | 184 | 0 | 0 | 10 | 194 |
| *ey* RNAi | 154 | 0 | 0 | 71 | 225 |
| *toy* RNAi | 190 | 14 | 0 | 67 | 271 |
| *sv* RNAi | 205 | 0 | 0 | 29 | 234 |
| *ey+toy* RNAi | 86 | 0 | 148 | 34 | 268 |
