## Supplementary material for "Pax6 homologs are required for patterning both visual systems of the daddy-longlegs *Phalangium opilio*": Table S5

**Table S5.** List of primers used for cloning and RNAi.

| **Gene** | **Forward Primer Sequence** | **Reverse Primer Sequence** |
| --- | --- | --- |
| *sv* | TAC TAC TCA TGT CAC GGC GG | GAG GTC ATC TTG TTT CGG CC |
| *ey* | GGC CGC GGC ACA GCG GCA TTA ACC AACT | CCC GGG GCC GTC TTG CGC TTG TAC TTGA |
| *toy* | GGC CGC GGA TGT CCC TCC ATC TTT GCCT | CCC GGG GCT TAA CTT CTC CTC CCG TCGC |
