## Supplementary figures and images for "Pax6 homologs are required for patterning both visual systems of the daddy-longlegs *Phalangium opilio*"

### Figure S1

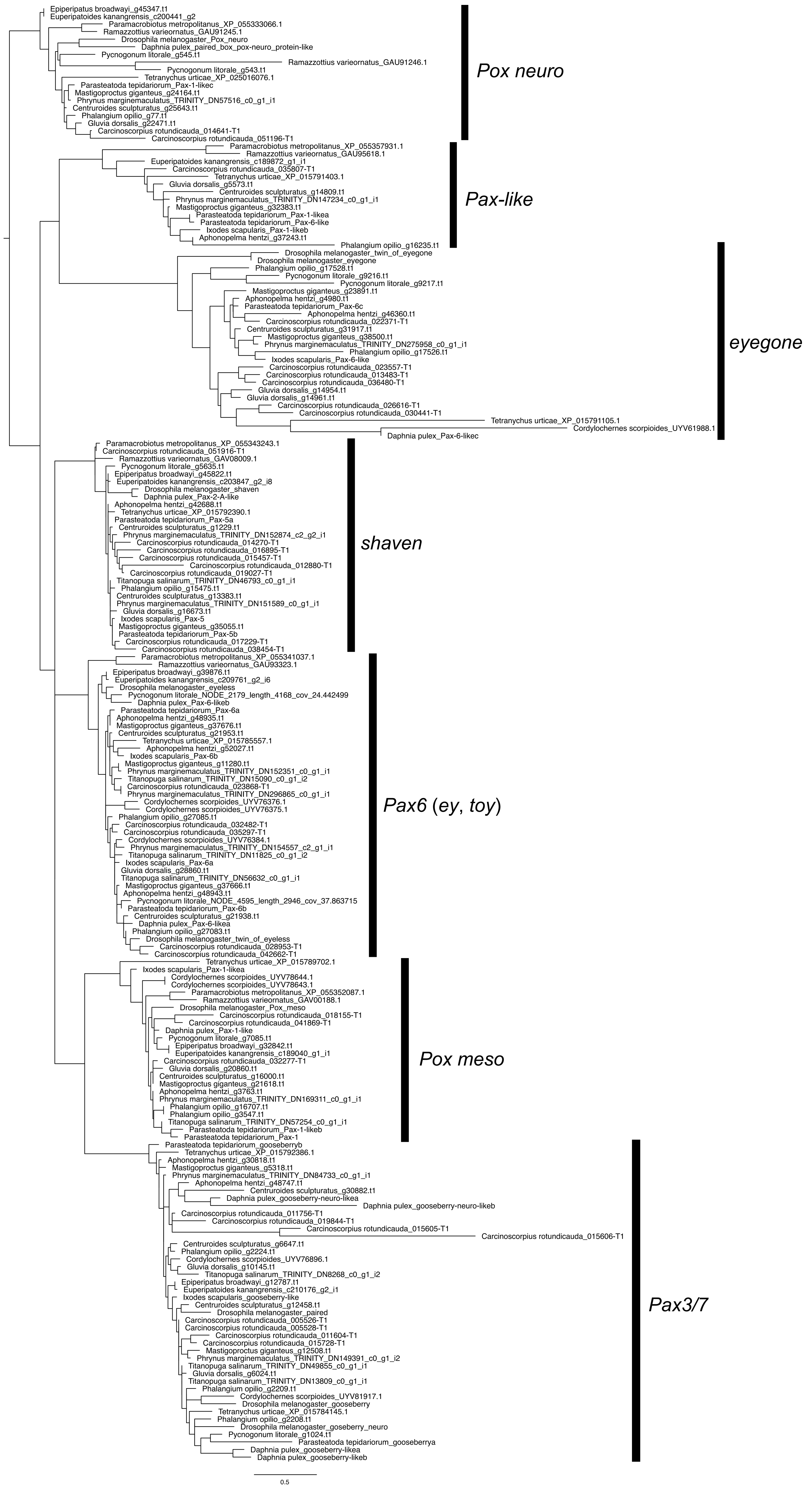

### Figure S2

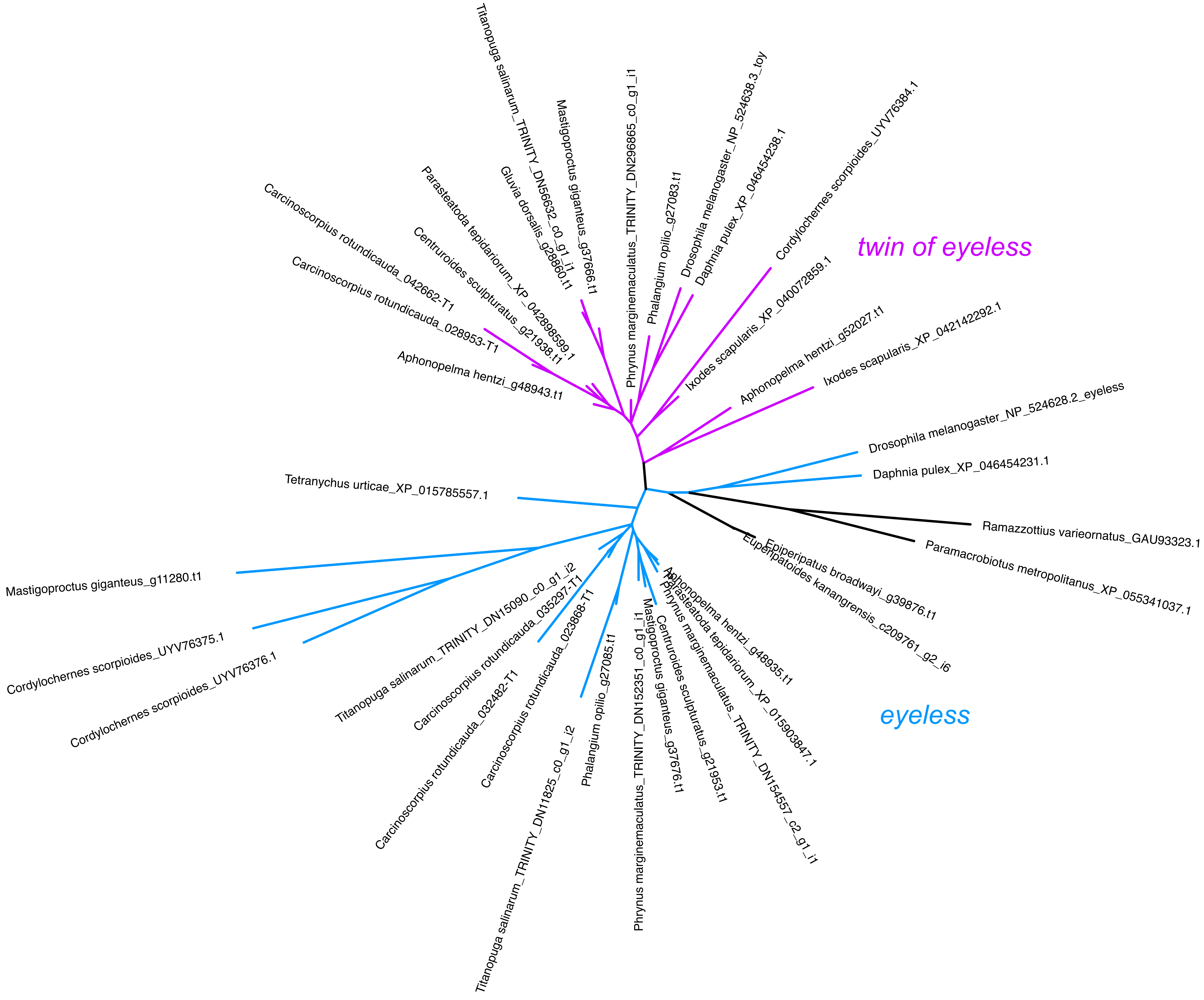

### Figure S3

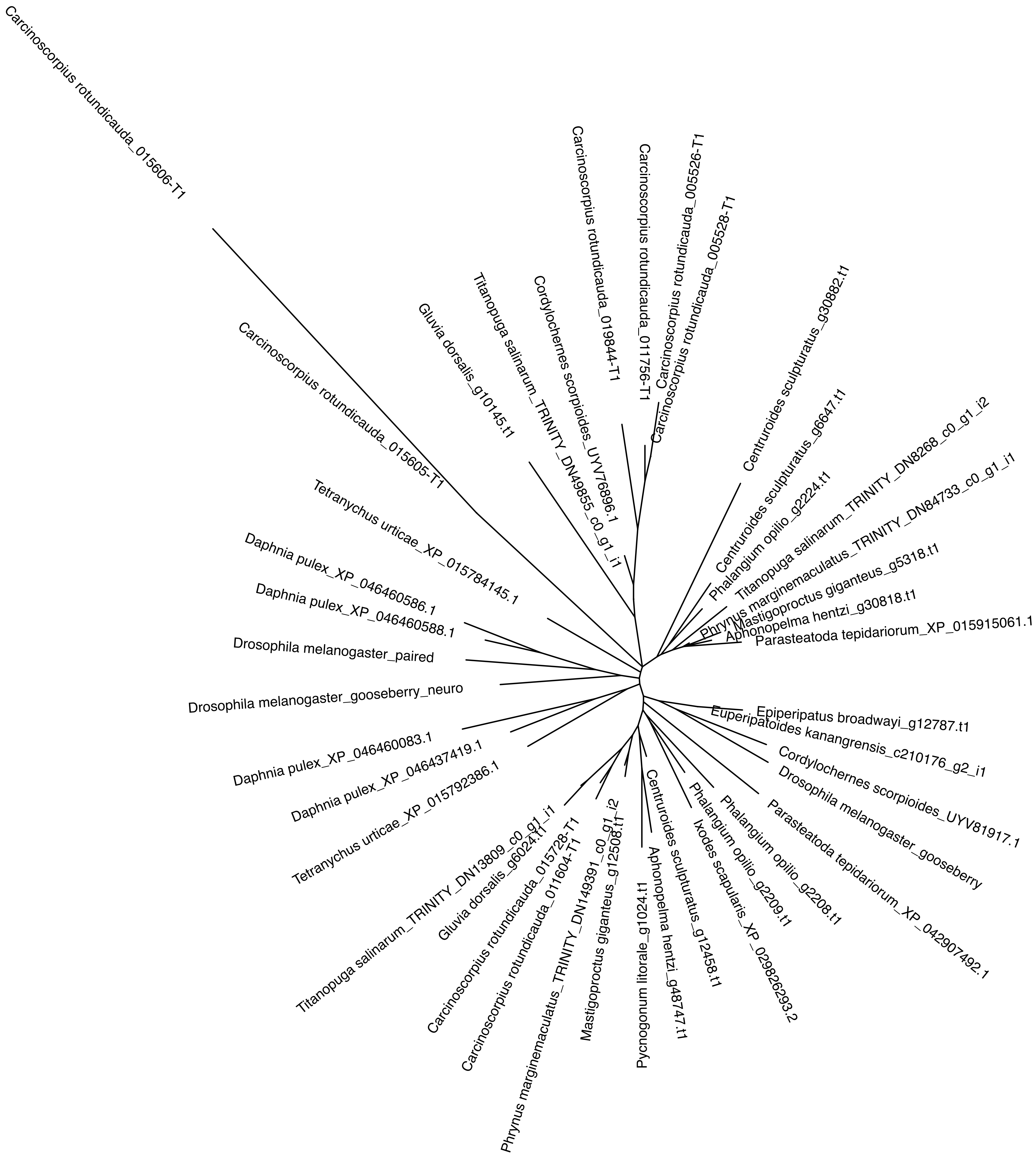

### Figure S4

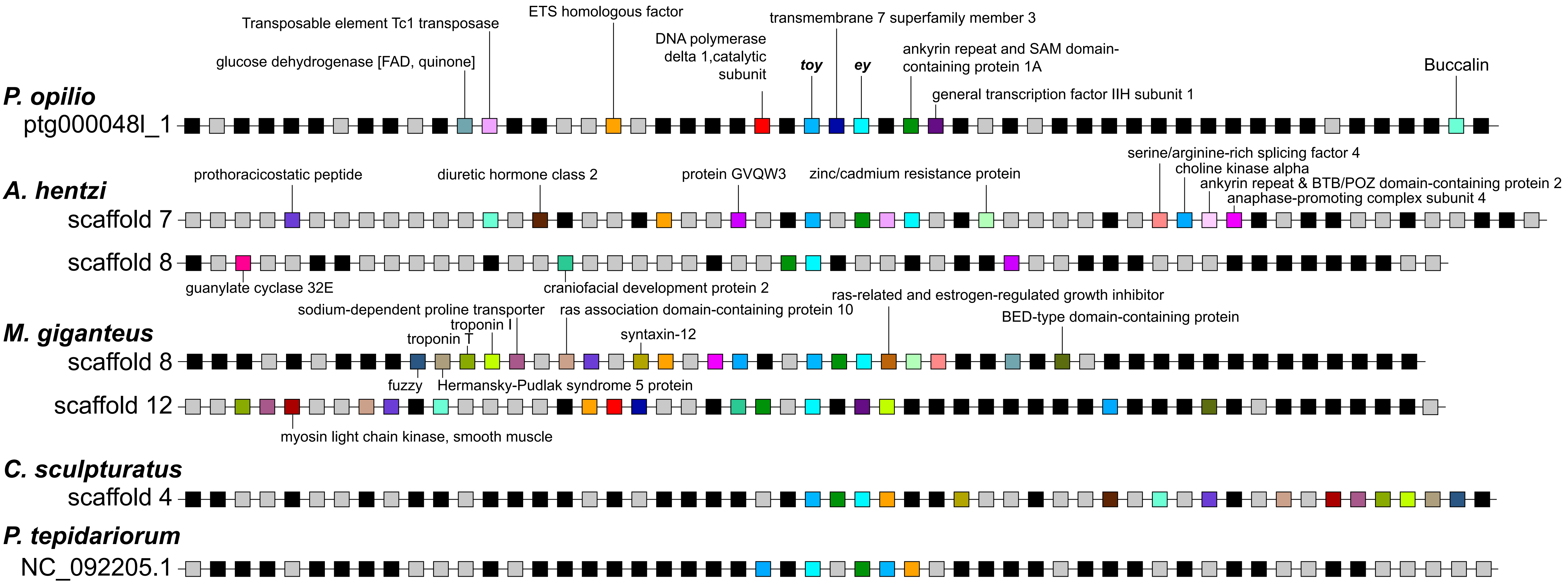

### Figure S5

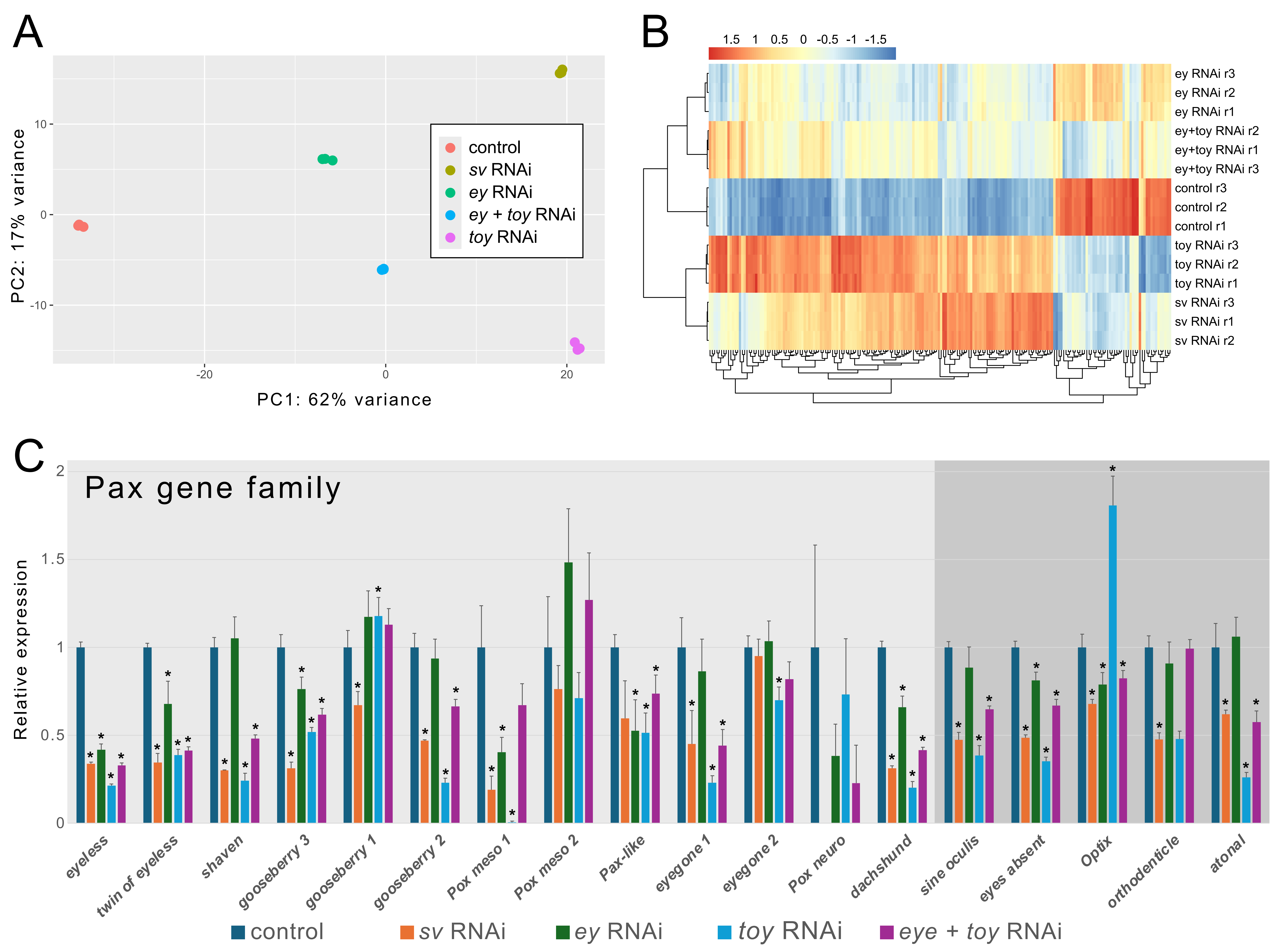

### Figure S6

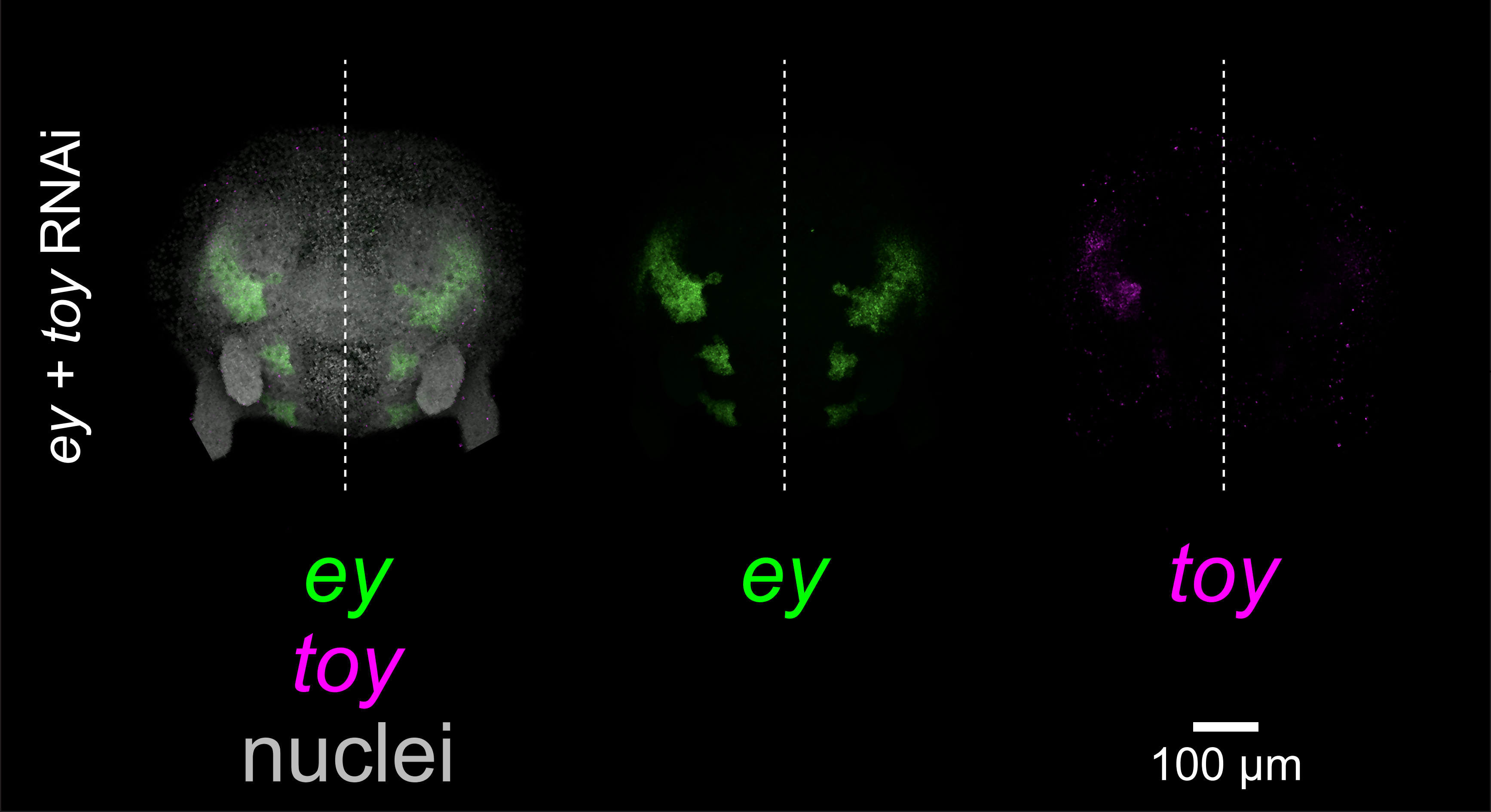
